# Population structure-guided profiling of antibiotic resistance patterns in clinical *Listeria monocytogenes* isolates from Germany identifies *pbpB3* alleles associated with low levels of cephalosporin resistance

**DOI:** 10.1101/2020.05.25.114330

**Authors:** Martin A. Fischer, Sabrina Wamp, Angelika Fruth, Franz Allerberger, Antje Flieger, Sven Halbedel

## Abstract

Case numbers of listeriosis have been increasing in Germany and the European Union during the last decade. In addition reports on the occurrence of antibiotic resistance in *Listeria monocytogenes* in clinical and environmental isolates are accumulating. The susceptibility towards 14 antibiotics was tested in a selection of clinical *L. monocytogenes* isolates to get a more precise picture of the development and manifestation of antibiotic resistance in the *L. monocytogenes* population. Based on the population structure determined by core genome multi locus sequence typing (cgMLST) 544 out of 1,220 sequenced strains collected in Germany between 2009 and 2019 were selected to cover the phylogenetic diversity observed in the clinical *L. monocytogenes* population. All isolates tested were susceptible towards ampicillin, penicillin and co-trimoxazole - the most relevant antibiotics in the treatment of listeriosis. Resistance to daptomycin and ciprofloxacin was observed in 493 (91%) and in 71 (13%) of 544 isolates, respectively. While all tested strains showed resistance towards ceftriaxone, the minimal inhibitory concentrations (MIC) observed varied widely between 4 mg/L up to >128 mg/L. An allelic variation of the penicillin binding protein gene *pbpB3* could be identified as the cause of this difference in ceftriaxone resistance levels. This study is the first population structure-guided analysis of antimicrobial resistance in recent clinical isolates and confirms the importance of penicillin binding protein B3 (PBP B3) for the high level of intrinsic cephalosporin resistance of *L. monocytogenes* on a population-wide scale.

## Introduction

*Listeria monocytogenes* is an important foodborne pathogen and the causative agent of listeriosis, an illness with symptoms ranging from gastroenteritis to septicemia, meningoencephalitis and miscarriage in pregnant women. *L. monocytogenes* infections are mostly associated with milk products, but also with meat, fish and vegetables [1]. Case numbers of listeriosis have been increasing during the last years with 701 notified cases in 2018 in Germany [2]. The incidence of listeriosis is relatively low (0.1-1.6 per 100,000 persons) compared to other gastrointestinal infections. However, fatality rates range between 7 to 30% despite antibiotic treatment [3,4]; even though *L. monocytogenes* is susceptible to a variety of antibiotics *in vitro*, it is one of the most fatal gastrointestinal foodborne bacterial pathogens.

The incubation period of listeriosis ranges from 1-67 days [5]. This rather long time frame complicates back-tracing of food vehicles through patient interviews and thus often has hampered the identification of outbreak sources. Whole genome sequencing (WGS)-based subtyping techniques, such as core genome multi locus sequence typing, have been implemented recently in many countries to improve disease cluster recognition and compare clinical and food isolates. This has enormously facilitated the identification of infection sources of listeriosis outbreaks [6–11].

The standard therapy for listeriosis is ampicillin or penicillin, frequently combined with gentamicin because a pronounced synergism between these antibiotics has been observed *in vitro* [12,13]. However, the effectivity of the combination therapy has been questioned in retrospective studies investigating the outcome of listeriosis treated either with the combination of both antibiotics versus penicillin monotherapy, with no benefit of the combined treatment on the patient’s outcome [14,15]. As an alternative, treatment with trimethoprim/sulfamethoxazole (hereinafter referred as co-trimoxazole) has been applied successfully in patients allergic to β-lactam antibiotics [16]. Meropenem is occasionally applied in listeriosis treatment, but therapy failure and mortality rate is higher under these conditions [17,18].

Antibiotic resistance to the clinically used antibiotics is rare in clinical isolates of *L. monocytogenes*, but recent studies provide evidence of increasing numbers of environmental isolates, including samples from animals, food and food-processing plants, with antibiotic resistance [19–21]. This observation is alarming since there is evidence that the increase of minimal inhibitory concentrations (MIC) observed in environmental strains also manifested in clinical strains later on [22]. Therefore, monitoring the development of antibiotic resistance in clinical isolates is of utmost importance to ensure appropriate antibiotic therapy of listeriosis in the future.

Beside the potential emergence of resistance to antibiotics used in standard therapy, *L. monocytogenes* is intrinsically resistant to third-generation cephalosporins such as ceftriaxone [13,23], often used to treat bacterial meningitis. Hence, as long as *L. monocytogenes* cannot be ruled out as the causative agent, co-administration of ceftriaxone or other cephalosporins with ampicillin is required [24]. Several factors including the penicillin binding protein PBP B3, encoded by the *lmo0441* gene, contribute to the intrinsic cephalosporin resistance of *L. monocytogenes* [25,26]. A *L. monocytogenes* mutant lacking *lmo0441* has strongly reduced cephalosporin resistance but did not reveal any other obvious phenotypes [25,27], suggesting that PBP B3 has a function specifically required during cephalosporin exposure.

Based on genome sequence data, we here designed a selection of 544 clinical *L. monocytogenes* strains covering the entire phylogenetic biodiversity observed among the strains isolated from human infections in Germany between 2009 and 2019. This strain selection was screened for antibiotic susceptibility against 14 clinically relevant antibiotics to describe the current antibiotic resistance levels of clinical *L. monocytogenes* strains on a population-wide scale. This led to the discovery of *pbpB3* mutations associated with reduced levels of cephalosporin resistance.

## Materials and Methods

### *L. monocytogenes* strains and growth conditions

All *L. monocytogenes* strains were grown on brain heart infusion (BHI) broth (# 211059, BD-BBL) or BHI agar plates (# CM0375, Oxoid) at 37°C. All strains used in this study are summarized in the supplementary Table S1.

### Construction of plasmids and strains

For expression of *pbpB3* variants in *L. monocytogenes, pbpB3* alleles of the strains 17-04405, 18-00287, 18-00792, 18-02573 and 18-04540 were amplified by PCR using the primers MF19 (5’-CGCGCCATGGATGGCTAGTTATGGTGGGAAAAAG) and MF20 (5’-CGCGGTCGACTTATTTATACATACTTTCAATAACTGGTTTTAGC). Fragments were cloned into plasmid pIMK3 [28] using NcoI/Sall. The sequence of the cloned inserts was confirmed by Sanger sequencing, the corresponding plasmid was introduced into strain LMJR41 (Δ*pbpB3*) by electroporation [28] and transformants selected on BHI agar plates containing 50 mg/L kanamycin. Correct plasmid insertion at the *attB* site of the tRNA^Arg^ was confirmed by PCR. The sequences of the above mentioned *pbpB3* alleles were submitted to NCBI GenBank (MT383155-MT383119).

### Genome sequencing

For genome sequencing, chromosomal DNA was extracted using the GenElute Bacterial Genomic DNA Kit (Sigma). One ng of the chromosomal DNA obtained was used in a library preparation using the Nextera XT library preparation kit (Illumina) according to manufacturer’s instructions. Sequencing was performed on Illumina MiSeq, NextSeq or HiSeq 1500 instruments, using either the MiSeq Reagent Kit v3 (600-cycle kit) or the HiSeq PE Rapid Cluster kit (version 2) in combination with an HiSeq Rapid SBS (version 2) sequencing kit (500-cycle PE or 150-cycle SE kit).

### Population structure analysis

Genome sequencing reads were assembled using the velvet algorithm. MLST sequence types (ST) and cgMLST complex types (CT) according to the seven housekeeping gene MLST scheme [29] and the 1,701 locus cgMLST scheme [6], respectively, were extracted from the assembled contigs by automated allele submission to the *L. monocytogenes* cgMLST server (http://www.cgmlst.org/ncs/schema/690488/). Clusters were defined as groups of strains with ≤10 different alleles between neighboring strains. Generation of the minimal spanning tree was performed in the “pairwise, ignore missing values” mode. All of the aforementioned steps were performed using the built-in functions of the Ridom^®^ SeqSphere Software package version 6.0.0 (2019/04).

### Antibiotic susceptibility testing

Antibiotic susceptibility testing was performed as a microdilution assay in accordance with the EUCAST guidelines in the January 2019 version [30]. Briefly, selected *L. monocytogenes* strains were streaked out on BHI agar plates and incubated at 37 °C for 24 h. Three to five colonies from each plate were picked, joined and further incubated in 3 mL BHI broth for 6 h. This culture was used to adjust NaCl solution (0.9%, w/w) to an OD_600_ of 0.005, representing a concentration of approximately 5·10^6^ colony forming units (CFU) per mL. Ten μl of this solution were used to inoculate the individual wells of a 96-well microtiter plate containing 90 μl Mueller-Hinton fastidious (MH-F) broth with the individual concentrations of the tested antibiotic; 1 mM IPTG was added where necessary. The overall plate design was adopted from a study by Noll and colleagues [21], produced in house and included ampicillin (AMP), benzylpenicillin (PEN), ceftriaxone (CRO), meropenem (MEP), daptomycin (DAP), ciprofloxacin (CIP), erythromycin (ERY), gentamicin (GEN), linezolid (LNZ), rifampicin (RAM), tetracycline (TET), tigecycline (TGC), vancomycin (VAN) and co-trimoxazole (SXT). Their concentrations were selected to cover the EUCAST-defined MIC breakpoints [30]. In cases where no breakpoint was defined for *L. monocytogenes*, the MIC breakpoints of *Streptococcus pneumoniae* or *Staphylococcus aureus* were used [30]. The plates were quickly mixed and incubated in a sealed polyethylene bag at 37 °C for 20±2 h. MIC were reported as the first concentration of the respective antibiotic where no visible growth was detected after the defined incubation period. A set of reference strains (*Escherichia coli* ATCC 259226, *Pseudomonas aeruginosa* ATCC 278538, *Staphylococcus aureus* ATCC 292139 and *Enterococcus faecalis* ATCC 29212) with known antibiotic resistance profiles were used to assure effectivity of the antibiotics under the chosen testing conditions.

### Association studies

The Kruskal-Wallis rank sum test was performed to determine if there were significant differences between samples in serogroups IIa, IIb and IVb as well as between different sequence types (where more than three strains were available) regarding the MIC values observed. To further test which groups significantly differed from one another, the pairwise Mann-Whitney-U test was performed. Adjusted p-values were obtained using a Bonferroni-Holm correction. All statistical analysis was performed using the stats package in R version 3.6.1 [31].

### Identification of alleles associated with reduced ceftriaxone resistance

Group-specific single nucleotide variations (SNV) were sought using the SNV tool implemented in SeqSphere (Ridom^®^, Germany). For this purpose, isolates with reduced ceftriaxone resistance belonging to a particular ST were defined as target and isolates outside this phylogenetic group as non-target. Moreover, isolates belonging to one of the other low-ceftriaxone resistance STs were excluded from the non-target group to increase sensitivity. SNVs occurring in 100% of the target group and which were different to 99% of the non-target group were accepted and only SNVs leading to non-synonymous amino acid exchanges were considered for further analysis.

## Results

### Population structure-guided isolate selection

The collection of clinical *L. monocytogenes* strains from the German consultant laboratory was used as the source of genetic diversity within the *L. monocytogenes* population. At the time this project was started, the collection contained 1,220 genome sequenced *L. monocytogenes* strains, isolated from human infections in Germany between 2009 and 2019. Of these strains 1,004 had been isolated from blood or cerebrospinal fluid and the remaining strains from other sources. Therefore, the majority of the strains (82%) were associated with invasive disease. Most of the strains were collected in 2016 (n=266), 2017 (n=395) and 2018 (n=453) (Figure S1).

The population structure of this strain collection was determined using MLST and cgMLST [6,11], allowing identification of disease clusters and sporadic cases. Most strains belonged to phylogenetic lineage I (57%, n=698) and lineage II (43%, n=520); cgMLST grouped the 1,220 isolates into 122 cgMLST complexes containing 798 isolates and 422 singletons. The 122 complexes varied in size from at least two up to 104 isolates, with a median size of 3 per complex (Figure S2). In order to cover all *L. monocytogenes* subtypes with current clinical relevance comprehensively, the following selection strategy was applied: At least one representative strain from each of the 122 identified complexes was selected. In cases where more than two isolates belonged to a cluster, its most central isolate was chosen. If strains with different CTs formed a joined complex, a representative strain belonging to the most abundant CT within this complex was selected. All sporadic isolates (422 of 1,220) were additionally included to further increase the genetic diversity within the selection of *L. monocytogenes* isolates. This procedure led to a selection of 544 *L. monocytogenes* strains from 2009 to 2019 with the majority of strains from 2016 to 2019 including representatives of the molecular serogroups IIa (39.7%), IIb (10.8%), IIc (1.3%), IVa (0.2%), IVb (46.7%), IVb-v1 (0.7%) and IVc (0.2%, Figure S1). Representatives of all 62 STs in the original strain collection were also present in this selection, with ST1, ST6 and ST2 representing the three most abundant STs (Figure S2B). Of the 587 CTs identified in the original strain collection, 539 (92%) were also included. Thus, the strain selection for antibiotic profiling contained 544 *L. monocytogenes* isolates in total and represents a miniaturized model collection of the clinical *L. monocytogenes* population currently causing infections in Germany (Table S1, Figure S1).

### Antibiotic profiling of the miniaturized model population

Each strain of the model population was tested for resistance against 14 clinically relevant antibiotics. No resistance was observed against ampicillin, penicillin or co-trimoxazole, which are the antibiotics currently recommended for the treatment of listeriosis (Table 1, Figure 1A). Still, two of the tested strains were susceptible to increased concentrations of ampicillin and penicillin and three isolates were susceptible to increased concentrations of co-trimoxazole. Among all strains tested one showed resistance to gentamicin, but genetic determinants explaining this phenotype were not identified. No resistance was observed to erythromycin, linezolid, meropenem, rifampicin, tigecycline and vancomycin. Furthermore, all isolates tested (544/544, 100%) were resistant to ceftriaxone (Table 1). This observation is in full agreement with the intrinsic cephalosporin resistance of *L. monocytogenes*. Moreover, the majority of the screened strains also showed resistance to daptomycin (493/544, 91%), a cyclic lipopeptide antibiotic. Around 13% of the isolates (71/544) showed resistance against the gyrase inhibitor ciprofloxacin. One strain was found to be resistant against tetracycline, against which most of the strains showed intermediate resistance (518/544, 95%). Susceptibility to increased concentrations was also observed for most isolates in case of linezolid (515/544, 95%) and ciprofloxacin (451/544, 83%), while it was less common with vancomycin (203/544, 37%), gentamicin (55/544, 10%), daptomycin (45/544, 8%) and meropenem (17/544, 3%). Sixteen strains showed growth in the presence of 0.6125 mg/L rifampicin, the lowest tested concentration, and must thus be considered as susceptible to increased doses.

**Figure 1:**
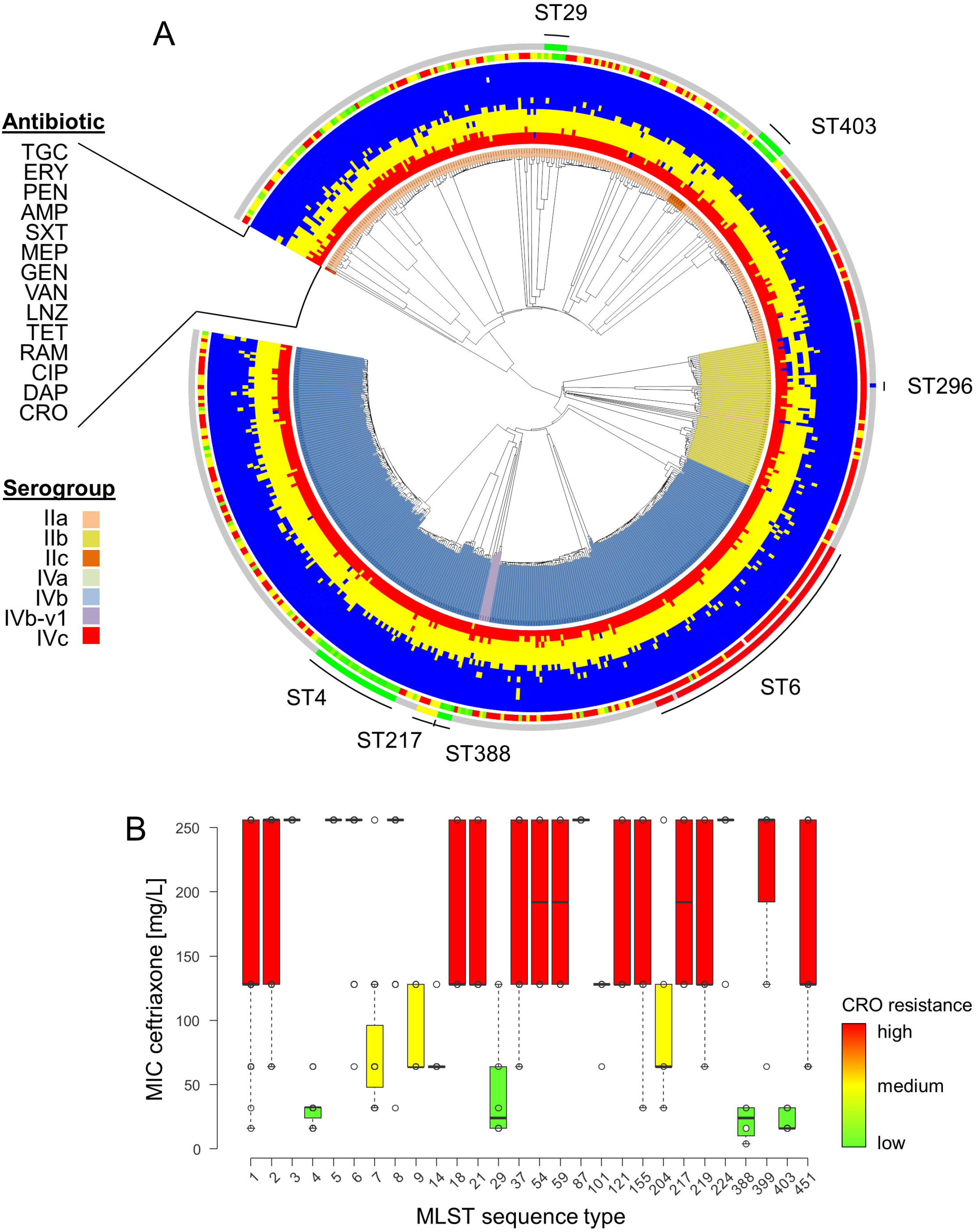
Identification of phylogenetic groups with reduced ceftriaxone resistance. (A) Phylogeny of isolates shown as Neighborhood-Joining tree based on the 1,701 locus cgMLST scheme for the model population used for the antibiotic susceptibility testing. Starting from the center, the rings represent the serogroups and the antibiotics tested (CRO, DAP, CIP, RAM, TET, LNZ, VAN, GEN, MEP, SXT, AMP, PEN, ERY, TGC). The color code for the antibiotics represent resistant (red), intermediate susceptible (yellow) and susceptible (blue) strains for the individual antibiotics. The two outer rings show MIC values determined for CRO from 4 mg/L (green) to >128 mg/L (red), as well as the positions of the isolates belonging to the STs further investigated. Data was visualized using iTOL v4 [51]. (B) Ceftriaxone resistance levels among 544 selected *L. monocytogenes* isolates according to MLST STs. Only STs for which MICs of ≥4 isolates were available were considered in this analysis.

**Table 1:**
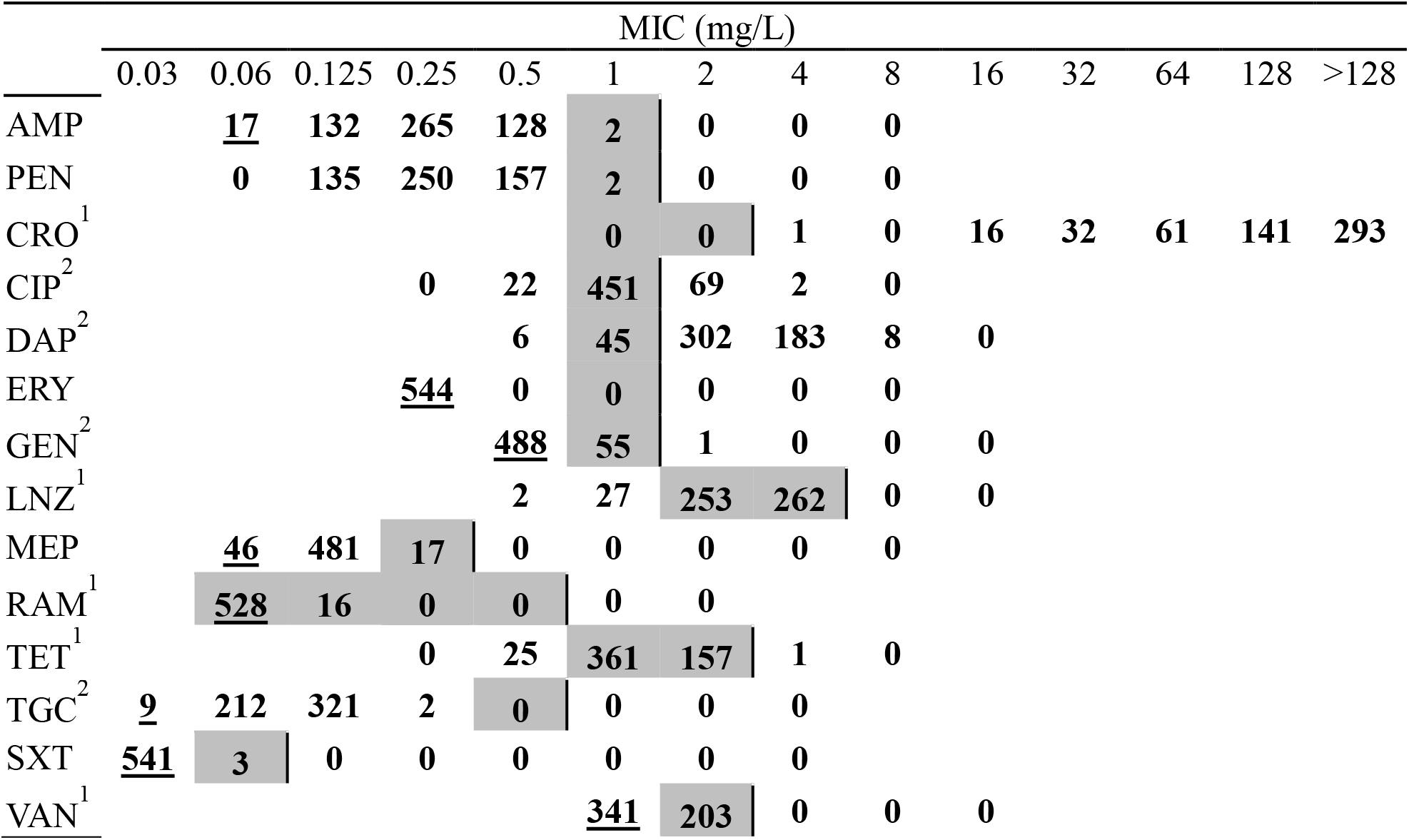
Antibiotic resistance profiles of the *L. monocytogenes* model population. 544 clinical *L. monocytogenes* strains were tested against 14 antibiotics. <u>Underlined</u> values indicate no observable growth at the lowest tested concentration. Concentrations in grey areas were not tested. Vertical lines indicate resistance breakpoints as defined by EUCAST for *Listeria monocytogenes, Streptococcus pneumoniae*^1^ or *Staphylococcus aureus*^2^. Intermediate resistance is marked by a grey background. All values below the grey area are considered fully susceptible, all values right of the vertical bar are considered fully resistant

The most common co-occurrence of antibiotic resistance was observed with ceftriaxone in addition to daptomycin (493/544, 91%). Out of these isolates, 66 (12%) showed additional resistance to ciprofloxacin. Only two isolates were found to be resistant to ceftriaxone and ciprofloxacin while being susceptible to daptomycin. Forty-five isolates (8%) were only resistant to ceftriaxone but none of the other antibiotics tested. Thus, they only showed intrinsic resistance against cephalosporins.

### Identification of phylogenetic groups with different antibiotic resistance profiles

The majority of isolates within the model population belonged to the molecular serogroups IIa, IIb and IVb (529 of 544, 97%). On the binary observation level of resistant versus sensitive, resistances were equally distributed between these three main molecular serogroups. To increase the resolution, the MIC values for each antibiotic were compared between isolates belonging to the different molecular serogroups. The average MICs for ampicillin (IVb=0.36 mg/L, IIa=0.18 mg/L), penicillin (IVb=0.38 mg/L, IIa=0.19 mg/L), daptomycin (IVb=2.94 mg/L, IIa=2.46 mg/L), linezolid (IVb=3.78 mg/L, IIa=2.13 mg/L), tetracycline (IVb=1.49 mg/L, IIa=1.11 mg/L), tigecycline (IVb=0.12 mg/L, IIa=0.08 mg/L) were significantly higher (Mann-Whitney U Test, n_1_=216, n_2_=254, P<0.05) for serogroup IVb isolates compared to isolates of serogroup IIa (Figure S3).

Despite this observation, we also found that the MICs for ceftriaxone varied between 4 mg/L up to >128 mg/L, with a median MIC of >128 mg/L considering all tested isolates (Table 1). While this classifies all strains as ceftriaxone-resistant, reduced median MIC values for ceftriaxone of ≤32 mg/L were found for certain STs (Figure 1B). The largest phylogenetic group with lowered ceftriaxone resistance was ST4 (n=24 isolates), showing a reduced median MIC of 32 mg/L in contrast to >128 mg/L for the remaining population. Likewise, lowered ceftriaxone MICs were observed for ST29 (median MIC=24 mg/L, n=7), ST388 (median MIC=24 mg/L, n=4) and ST403 isolates (median MIC=16 mg/L, n=8, Figure 1B).

Reduced ceftriaxone resistance levels were also observed in ST7 (median MIC=64 mg/L), ST9 (median MIC=64 mg/L), ST101 (median MIC=128 mg/L) and ST204 isolates (median MIC=64 mg/L).

### Identifying *pbpB3* alleles linked to reduced ceftriaxone resistance

Single nucleotide variant analysis revealed that ST4, ST29, ST388 and ST403 isolates associated with lowered levels of ceftriaxone resistance carried group-specific non-synonymous mutations in various coding regions. However, the only gene carrying one mutation common to all isolates belonging to the STs with reduced ceftriaxone resistance was *lmo0441*, encoding PBP B3, which showed a mutation within the allelic version found in ST4 and ST388 (*pbpB3* allele type 4, Ala172Val) and ST403 and ST29 (*pbpB3* allele type 20, Thr53Ser, Figure 2A,B). This suggests that certain *pbpB3* alleles are associated with reduced resistance against ceftriaxone. Remarkably, all ST4 and ST388 isolates carried the *pbpB3 Ala172Val* substitution characteristic for *pbpB3* allele no. 4 in the Ruppitsch cgMLST scheme and this *pbpB3* allele was not found in any other strain. Likewise, all our ST403 isolates carried the *pbpB3 Thr53Ser* variant (allele no. 20), also found in four out of six ST29 isolates tested with lowered ceftriaxone resistance levels. The two ST29 isolates tested with a ceftriaxone resistance above the median value observed in this group had a different *pbpB3* allele. Despite its presence in these two subgroups, *pbpB3* allele no. 20 was not found in any other of the 1,220 strains of the original strain collection. We thus conclude that *pbpB3* alleles 4 and 20 are associated with reduced ceftriaxone resistance.

**Figure 2:**
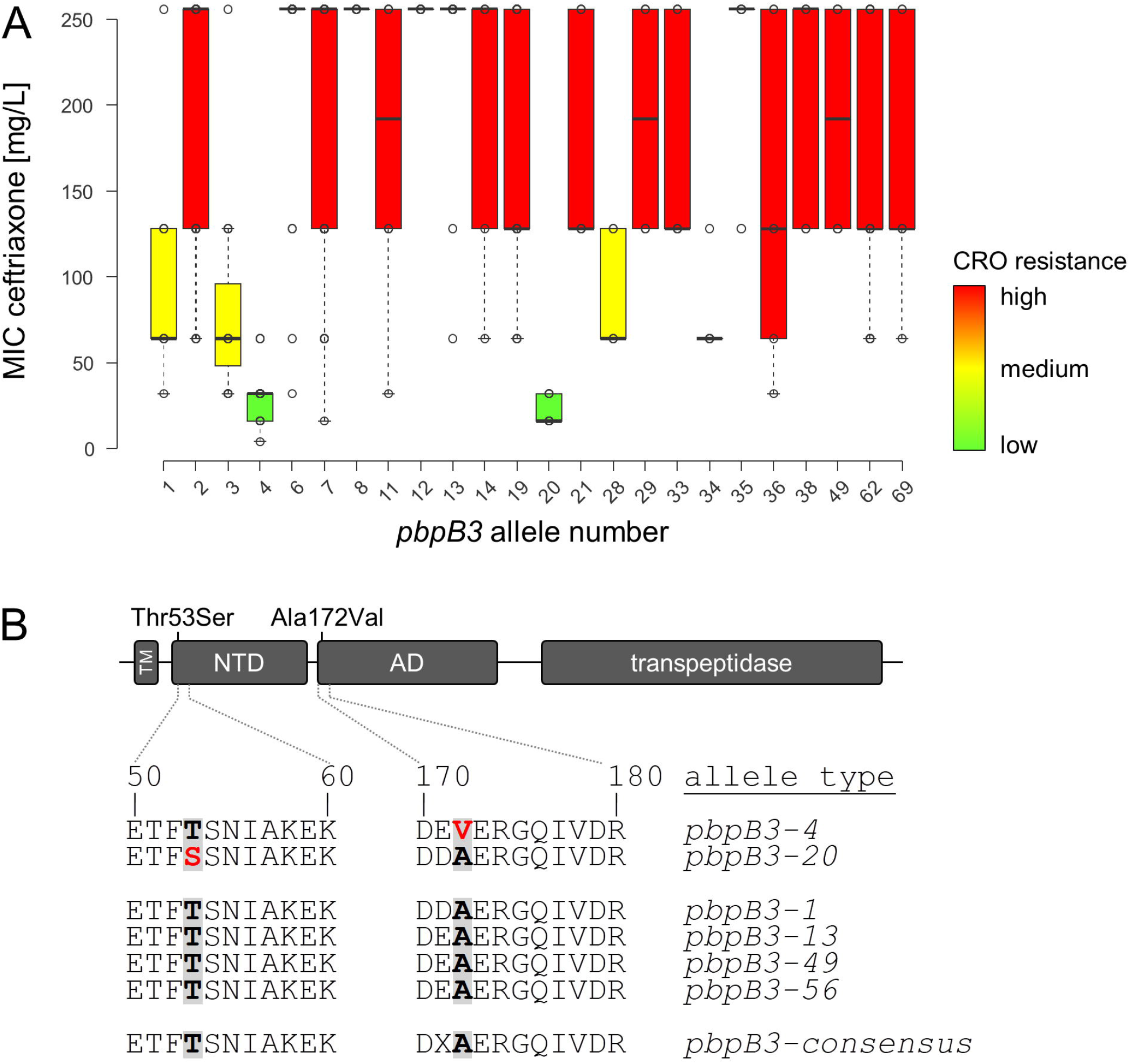
Identification of *pbpB3* alleles associated with reduced ceftriaxone resistance. (A) Ceftriaxone resistance levels among 544 selected *L. monocytogenes* isolates according to their *pbpB3* allele in the Ruppitsch cgMLST scheme. Only those *pbpB3* alleles for which MICs of ≥4 isolates were available were considered in this analysis. (B) Scheme illustrating PBP B3 domains and position of the amino acid exchanges found in the *pbpB3* alleles no. 4 (Ala172Val) and 20 (Thr53Ser), which are associated with reduced ceftriaxone resistance. Abbreviations: TM - transmembrane helix, NTD - N-terminal domain; AD - allosteric domain.

### Effect of novel *pbpB3* mutations on ceftriaxone resistance

Even though the sequence alterations in the two *pbpB3* alleles were rather conservative at the protein level, their contribution to ceftriaxone resistance was tested in a complementation assay. For this purpose, a Δ*pbpB3* deletion mutant constructed in the background of *L. monocytogenes* EGD-e (strain LMJR41) [27] was complemented with different *pbpB3* alleles and ceftriaxone resistance of the resulting strains was determined. In good agreement with previous results [25], ceftriaxone resistance was greatly reduced in the Δ*lmo0441* mutant (2 mg/L) compared to wild type strain EGD-e (64 mg/L). Reintroduction of the wild type *pbpB3* allele (allele type 1) from EGD-e restored this phenotype almost completely (32 mg/L). In contrast, expression of *pbpB3* allele type 4 associated with reduced ceftriaxone resistance in the Δ*pbpB3* background led to a lower ceftriaxone resistance level of only 16 mg/L. When *pbpB3* allele type 49, originating from a closely related but fully ceftriaxone-resistant ST217 isolate (MIC >128 mg/L, n=6), was expressed in the Δ*pbpB3* background, ceftriaxone resistance increased to 32 mg/L. This level of ceftriaxone resistance further increased to 64 mg/L, when *pbpB3* allele type 13 from ST6 strain 18-04540 - showing the highest observed level of ceftriaxone resistance in this study - was used for complementation. The complementation of the deletion mutant with *pbpB3* allele type 56 increased the ceftriaxone MIC to 32 mg/L. This allele type is identical to *pbpB3* allele type 4 except for a single mutation at the aforementioned position 172, where it still carries the original alanine. These results further underline the apparent importance of this single amino acid for the resistance against ceftriaxone. As for the *pbpB3* allele type 4, complementation mutants carrying *pbpB3* allele type 20 showed higher ceftriaxone compared to the deletion mutant but lower ceftriaxone resistance compared to the complementation mutants carrying *pbpB3* alleles of the wild type strain or from the high level resistance strain. In conclusion, *pbpB3* alleles from strains with low and high levels of ceftriaxone resistance confer low and high levels of ceftriaxone resistance upon their heterologous expression in the *ΔpbpB3* mutant, respectively. This confirms the association of certain *pbpB3* alleles with ceftriaxone resistance and demonstrates the population-wide validity of the concept that PBP B3 is an important determinant for ceftriaxone resistance in *L. monocytogenes*.

To estimate the overall relevance of this observation for the entire *L. monocytogenes* population, the frequency of *pbpB3* allele types 4 and 20 was calculated for the model population of 544 strains (55 unique *pbpB3* allele types), for the initially used clinical strain collection of 1,220 strains (58 unique *pbpB3* allele types) as well as for 27,118 *L. monocytogenes* genomes available on the National Center for Biotechnology Information (NCBI) pathogen detection pipeline at the time of this study (1,033 unique *pbpB3* allele types). Allele type 4 was detected in 28 strains of the model collection (expected: 10), in 39 strains of the clinical strain collection (expected: 21) and 340 times in the NCBI dataset (expected: 26). Allele type 20 was detected in 12 strains of the model collection, 62 strains of the clinical strain collection and 156 strains of the NCBI dataset. Therefore, the abundance of both allele types was above the theoretically expected values and hence the presence of theses *pbpB3* allele types does not seem to provide an evolutionary disadvantage.

## Discussion

Our results represent the first comprehensive determination of antibiotic resistance patterns of clinical *L. monocytogenes* strains isolated in Germany. The complexity of this strain collection was reduced by the generation of a non-redundant model population using cgMLST subtyping data. This model population contains less than half of the original isolates but still maintains the large biodiversity observed in the original *L. monocytogenes* clinical strain collection; determination of antibiotic resistance patterns in this model population greatly facilitated experimental determination of antibiotic resistance patterns without losing phylogenetic resolution.

An important finding of this study is the sustained effectivity of the standard antibiotics recommended for the treatment of listeriosis. None of the *L. monocytogenes* strains tested here showed full resistance against ampicillin and penicillin and only one was resistant towards gentamicin. However, gentamicin is not used as a stand-alone antibiotic in listeriosis therapy and only administered in combination with ampicillin or penicillin. Moreover, none of the isolates tested showed full resistance against co-trimoxazole, which is used as an alternative in patients with β-lactam allergy. However, susceptibility only to increased concentrations of penicillin (2/544), ampicillin (2/544) and co-trimoxazole (3/544) was observed in few cases. Therefore our results are in accordance with observations made with other clinical strain collections from Europe where intermediate resistance levels against these three antibiotics were also reported to occur with low frequency [22,32].

While the low level of resistance towards currently clinically applied antibiotics is a relief, the situation in environmental and food isolates is more alarming. *L. monocytogenes* strains with multi drug resistance or resistance to ampicillin, penicillin or co-trimoxazole have repeatedly been isolated from the environment and from different food types [19, 21, 33–38]. It can be expected that the antibiotic resistances observed in environmental and food strains today will later manifest in clinical strains. Therefore, surveillance of antimicrobial resistance development in clinical *L. monocytogenes* strains in the future is of great importance, especially since average resistance levels against several β-lactams have been continuously increasing since the 1920s in clinical *L. monocytogenes* isolates from France [22].

The highest level of resistance within our model population was observed for ceftriaxone (100%), to which *L. monocytogenes* is intrinsically resistant [13,23], daptomycin (91%) and ciprofloxacin (13%). However, breakpoints have not been established for daptomycin and ciprofloxacin in *L. monocytogenes* (as none of them is recommended to treat listeriosis) and applications of cephalosporins and ciprofloxacin have caused therapy failure in the past [39,40,41].

A large variation of ceftriaxone MICs ranging from 4 mg/L up to >128 mg/L was observed between isolates belonging to different STs and could be traced back to amino acid exchanges in *pbpB3*. Interestingly, an almost similar degree of variation in ceftriaxone resistance was observed within the ST1, ST155, ST451 strains included here (Figure 1B), even though no association between ceftriaxone resistance and *pbpB3* allele variation was found in these STs. Cephalosporin resistance is a multifactorial process in *L. monocytogenes* [26], and genetic variations in other cephalosporin resistance determinants may account for the variability of ceftriaxone resistance in these phylogenetic groups.

The PBP B3 of *L. monocytogenes* belongs to the same subclass of class B PBPs as *Bacillus subtilis* PBP3, *Staphylococcus aureus* PBP2a (encoded by *mecA*) and *Enterococcus faecalis* PBP5, which all are low-affinity penicillin binding proteins and as such critical determinants of cephalosporin or methicillin resistance in these bacteria [42–45]. The two *pbpB3* mutations lowering cephalosporin resistance described here affect the N-terminal domain and the allosteric domain (non-penicillin binding domain) of PBP B3 (Figure 2B). The function of these non-catalytic domains is not entirely clear, but amino acid exchanges in the allosteric domain of *S. aureus* PBP2a (such as N146K and E150K) are associated with increased resistance of *S. aureus* to ceftaroline, a fifth-generation cephalosporin [46–49]. Ceftaroline non-covalently interacts with this allosteric domain inducing a conformational change that makes the active site in the transpeptidase domain accessible for acylation and thus for inhibition by a second ceftaroline molecule [50]. The N146K and E150K mutations of *S. aureus* PBP2a map to the same stretch in the beginning of the allosteric domain as the A172V exchange in PBP B3 of *L. monocytogenes*. Apparently, amino acid exchanges in this region of the allosteric domain improve or impair cephalosporin binding in low affinity PBPs and thus resistance of different Gram-positive pathogens to this important group of antibiotics.

**Table 2:**
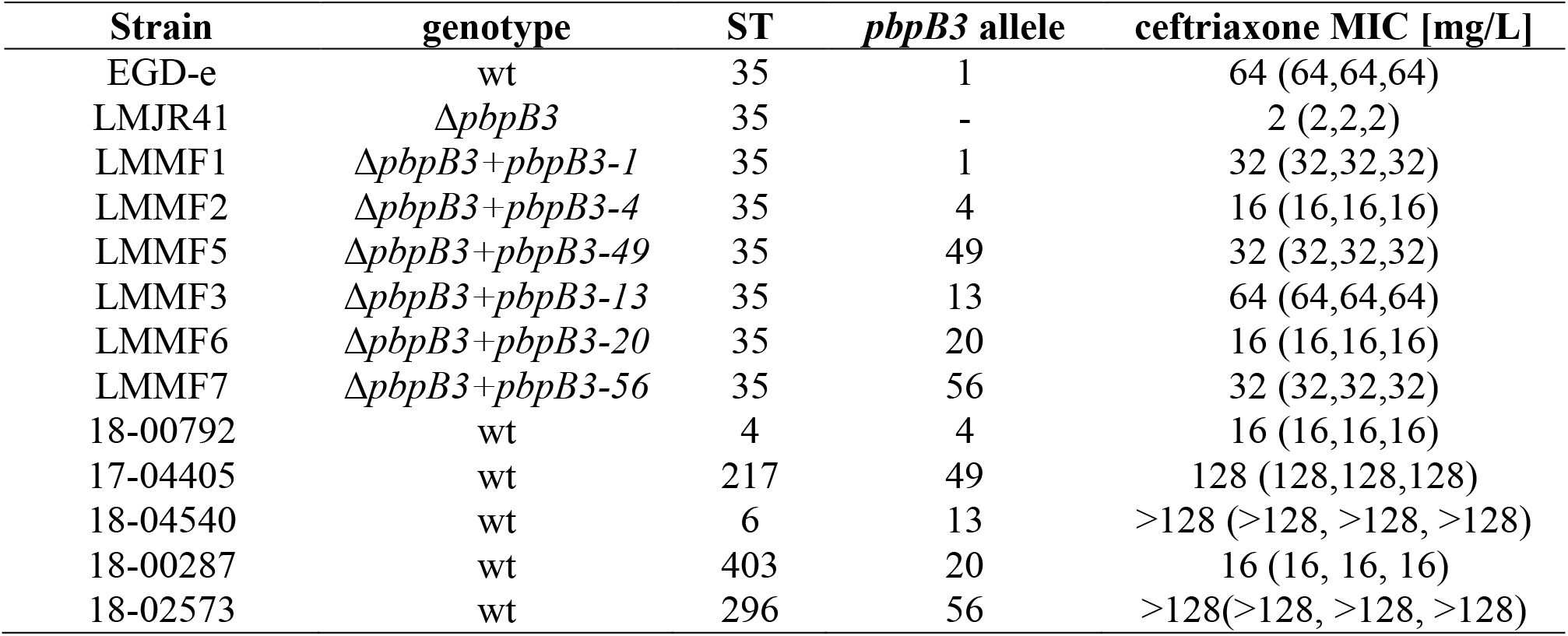
Effect of *pbpB3* on ceftriaxone resistance. MIC of ceftriaxone for *L. monocytogenes ΔpbpB3* strains complemented with *pbpB3* alleles from clinical *L. monocytogenes* strains with different levels of ceftriaxone resistance. All measurements were performed in triplicates and average values are shown with the individual values given in parentheses.

## Acknowledgements

The authors would like to acknowledge Matthias Noll and Ingo Klare for fruitful discussions and Karsten Großhennig, Petra Hahs and Claudia Lampel for technical assistance.

