## Supplementary Figures S1-S4 for "Population structure-guided profiling of antibiotic resistance patterns in clinical *Listeria monocytogenes* isolates from Germany identifies *pbpB3* alleles associated with low levels of cephalosporin resistance"

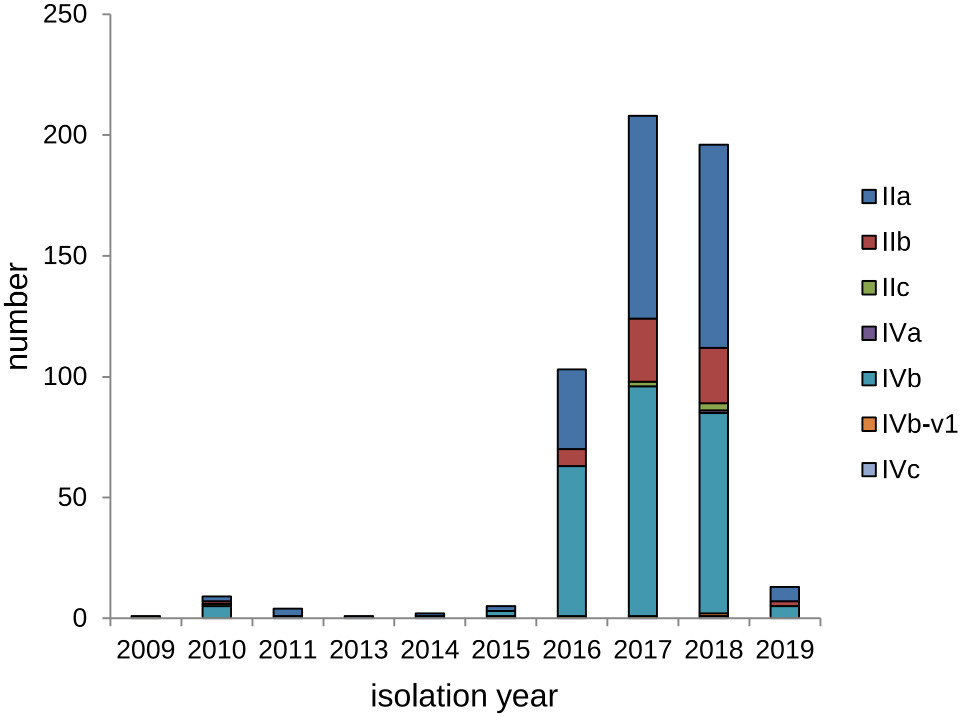


**Figure S1:** Isolation year in the selection of 544 clinical *L. monocytogenes* isolates that were subjected to antibiotic susceptibility testing.

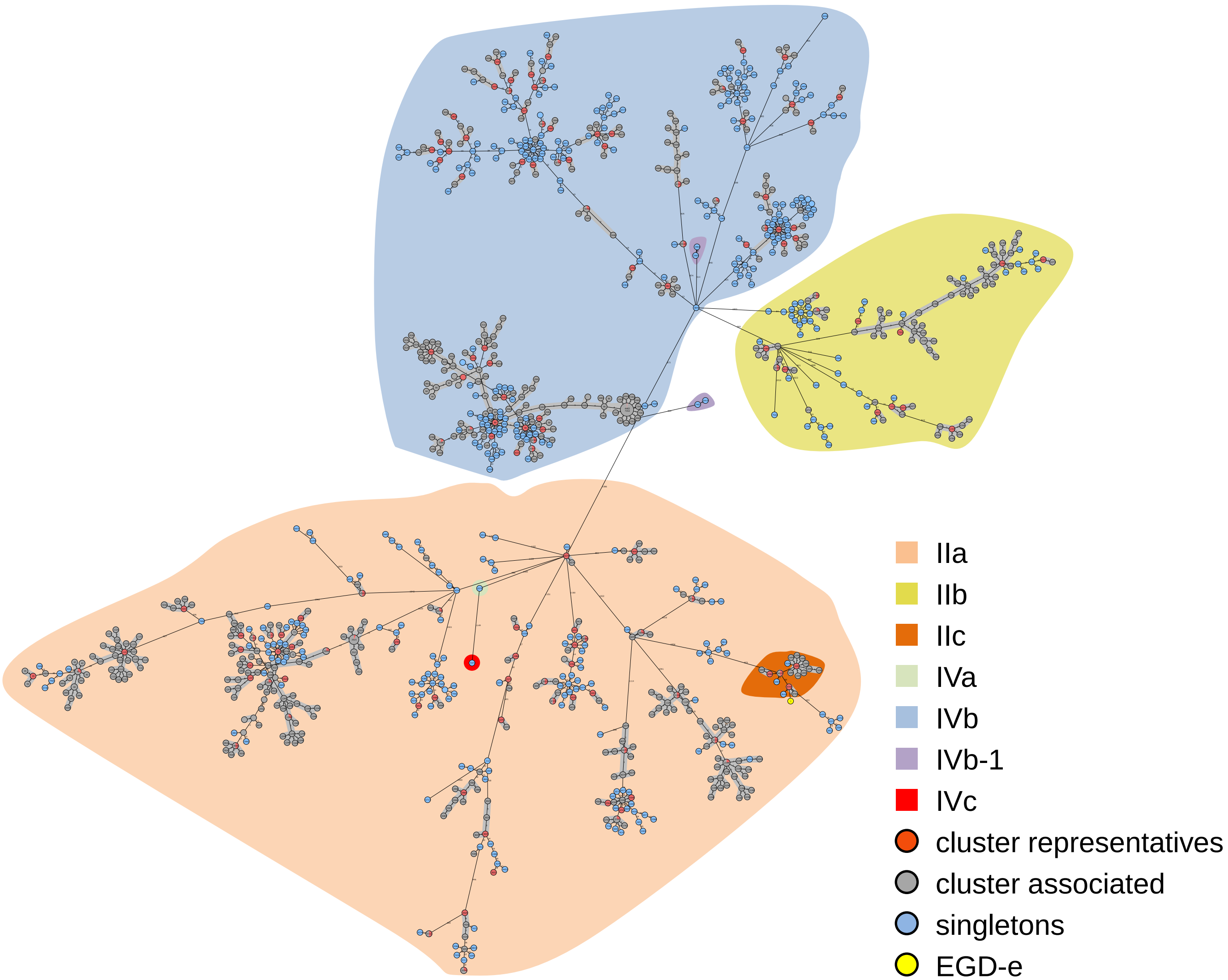


**Figure S2:** Minimum spanning tree of 1220 clinical *L. monocytogenes* isolates based on their cgMLST profiles. Samples screened for antimicrobial resistance patterns are colorized in red (outbreak cluster representatives) and blue (single isolates). The background is colorized according to molecular PCR serogroups.

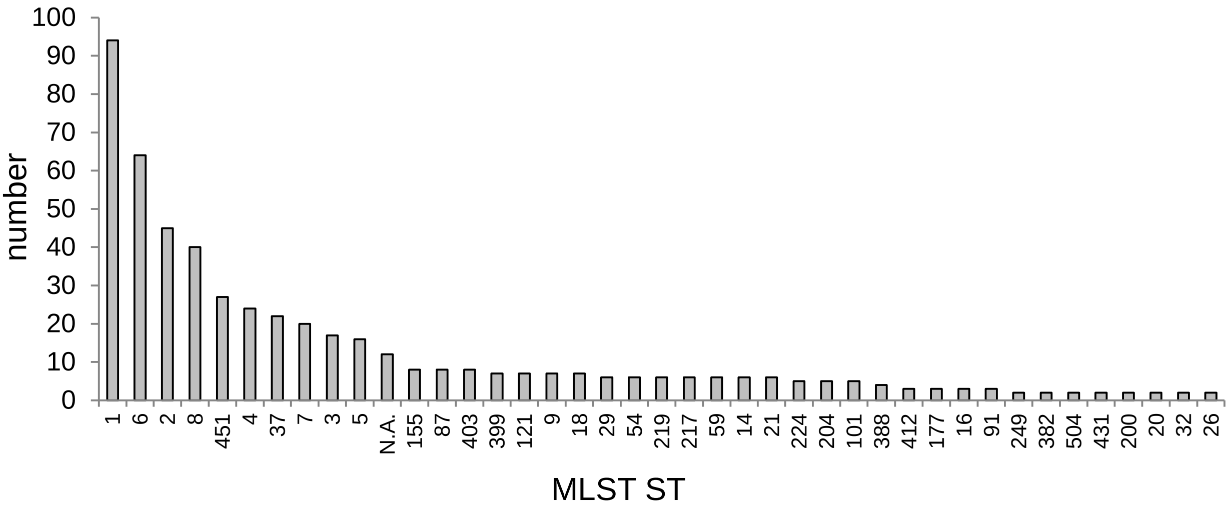


**Figure S3:** MLST sequence types (STs) in the selection of 544 clinical *L. monocytogenes* isolates that were subjected to antibiotic susceptibility testing. Only STs with more than one strain are shown.

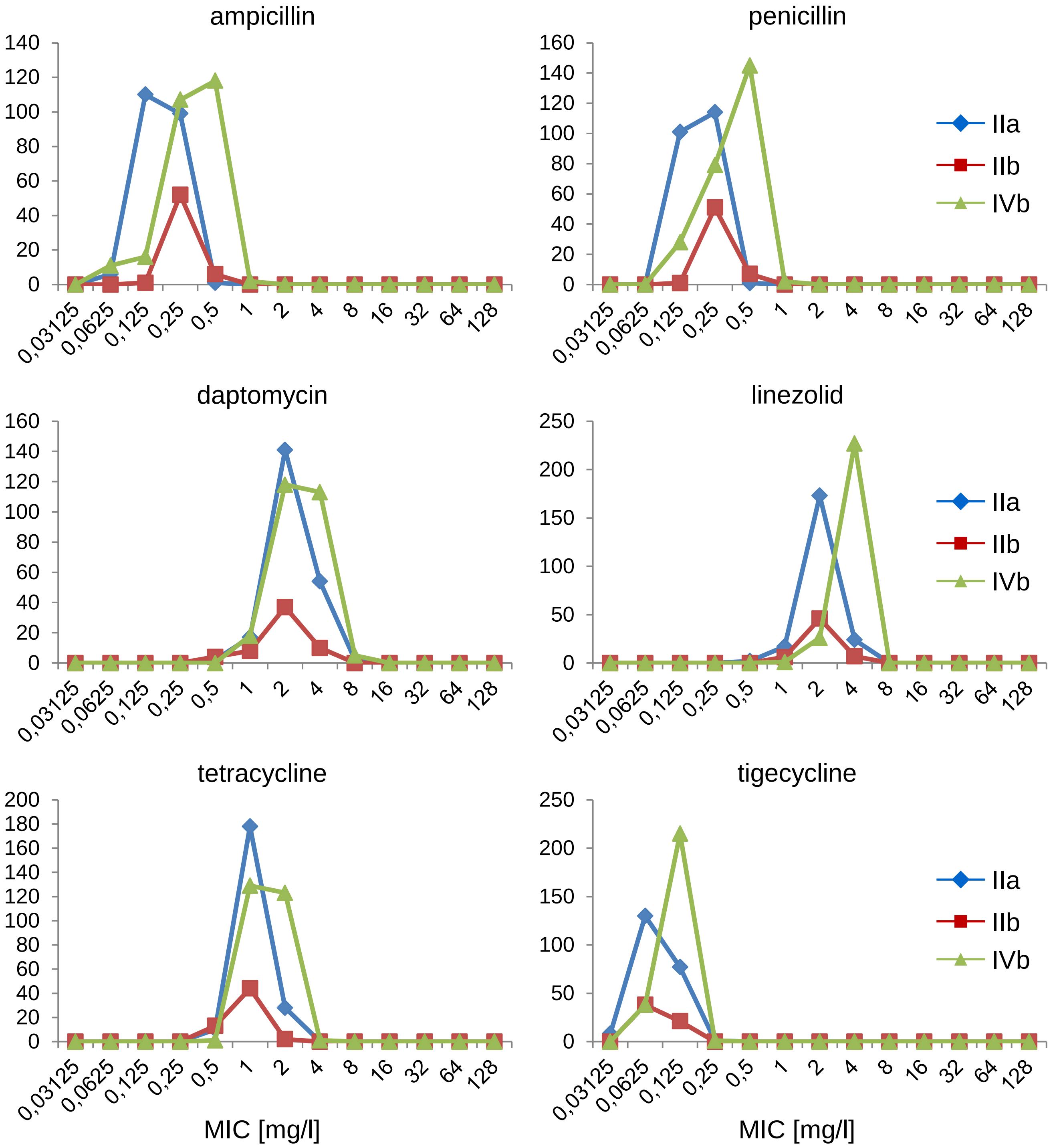


**Figure S4:** Distribution of minimal inhibitory concentrations for different antibiotics in *L. monocytogenes* isolates belonging to molecular PCR serogroups IIa (n=216), IIb (n=59) and IVb (n=254).
